# General and robust covalently linked graphene oxide affinity grids for high-resolution cryo-EM

**DOI:** 10.1101/657411

**Authors:** Feng Wang, Yanxin Liu, Zanlin Yu, Sam Li, Yifan Cheng, David A. Agard

## Abstract

Despite their great potential to facilitate rapid preparation of quite impure samples, affinity grids have not yet been widely employed in single particle cryo-EM. Here, we chemically functionalize graphene oxide coated grids and use a highly specific covalent affinity tag system. Importantly, our polyethylene glycol spacer keeps particles away from the air-water interface and graphene oxide surface, protecting them from denaturation or aggregation and permits high-resolution reconstructions of small particles.

## Main

Owing to a series of technological breakthroughs, cryo-electron microscopy (cryo-EM) is fast becoming the method of choice for determining protein structures at a near-atomic or atomic resolution^1, 2^. By contrast with the rapidity of data collection and processing, sample preparation remains slow and, in many cases, has become rate-limiting. Cryo-EM specimens are typically prepared by depositing purified proteins onto cryo-EM grids, which are usually a metal grid covered with continuous perforated carbon or gold films. After removing excessive sample solutions through blotting, the grid is plunged into liquid ethane and the biological samples are vitrified in amorphous ice^3^. Proteins of interest are thus preserved in hydrated state^4^. For challenging systems and dynamic complexes, the typical path of overexpression, biochemical-scale purification and concentration can be problematic, either because the system cannot be reconstituted, or aggregation may occur. Perhaps even more significant is the disruption of protein structure and protein-protein interactions that can occur when exposed to the air water interface^5^. During the formation of the very thin vitreous ice films (often <50nm) required for high resolution imaging, the very high surface area to volume ratio dramatically increases the probability of exposure to the denaturing interface. Indeed, most particles are observed at the air water interface^6^, and obtaining high resolution structures typically requires selecting only a small subset of picked particles. A potential solution to both the sample preparation and air-water interface problems is through the use of “affinity grids” that would simultaneous concentrate the sample on the grid while restricting it from the air-water interface. Over the past decades several affinity grid strategies have been proposed for cryoEM including decorating a supporting film with nickel-nitrilotriacetic acid (Ni-NTA) to capture His-tagged proteins^5, 7-9^, use of 2D streptavidin crystals to capture biotin or strep-tagged proteins^10^, or specific antibodies^11, 12^ to capture viruses.

Yet despite these promising advances, no single strategy has proven broadly useful –due to the inefficiency of His-tags, the high scattering of thick support films, the need for specific antibodies, etc. Here, we present a novel affinity grid approach that combines a small, essentially infinite affinity covalent tagging system with chemically derivatized graphene oxide (GO) support films only 1-2 molecules thick (**Fig 1a**). The self-ligating SpyCatcher/SpyTag coupling system^13^ forms a covalent bond between the ∼14KDa SpyCatcher and a 12 residue peptide, either of which can be fused to the protein of interest and the partner then coupled to the grid. The irreversible covalent bond forms within minutes^13^, is highly specific and robust, paving the way for “purification on the grid”. Graphene oxide (GO) was selected as the supporting film because (i) it significantly reduces background compared to amorphous carbon^14^, (ii) it is decorated with abundant oxygen-containing functional groups which facilitate further chemical modification, and (iii) is more straightforward to make and to coat grids than pure graphene crystals. Once a grid matrix and tagging system have been selected, the next challenge is to reduce non-specific grid interactions and to position the sample away from the grid surface as well as the air-water interface thereby avoiding potential sample denaturation or preferred orientation problems. For this, we choose to use a polyethylene glycol (PEG) spacer as it is well appreciated from single molecule studies that it is very effective in blocking non-specific adsorption.

**Figure 1:**
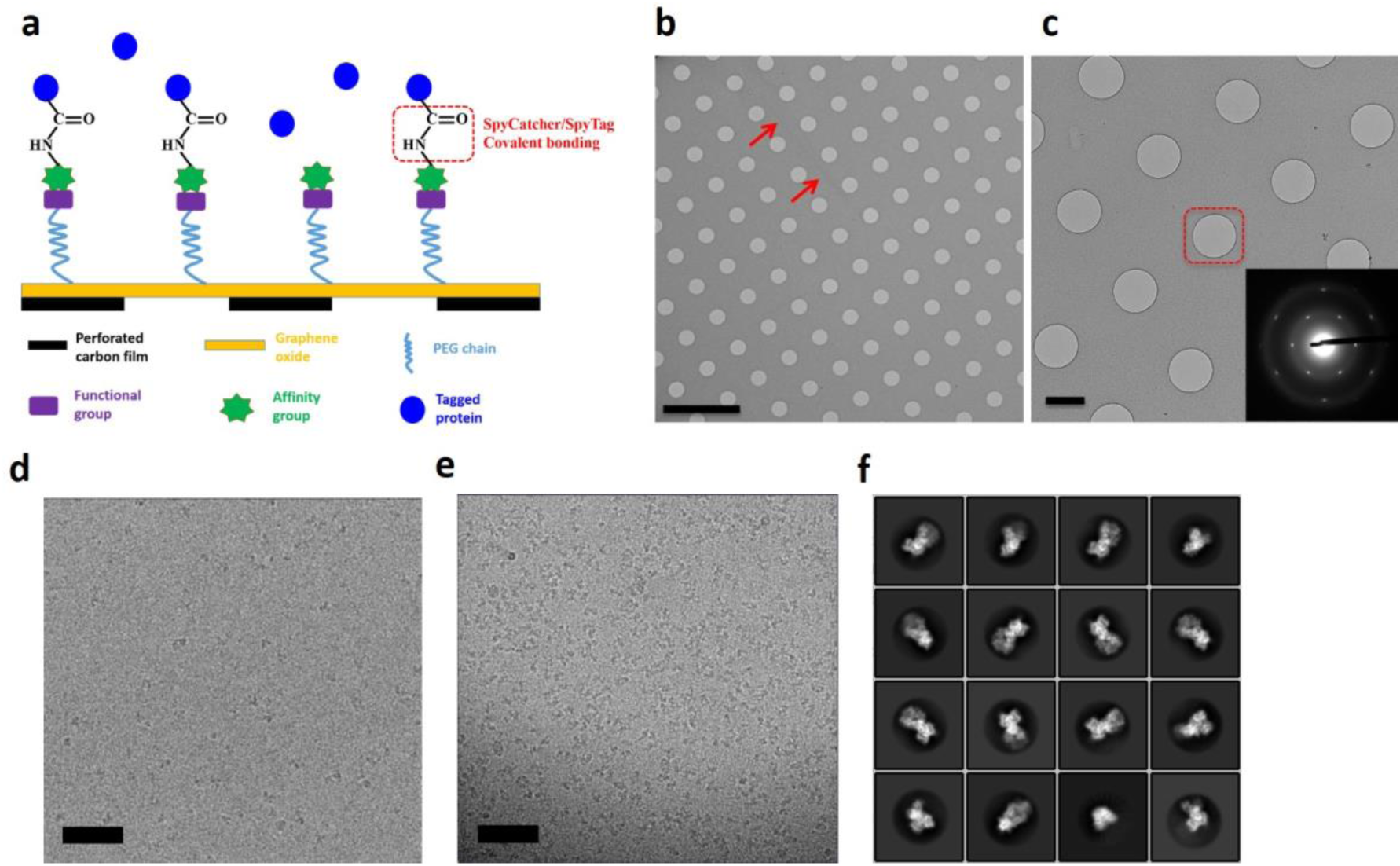
**a**, Schematic illustration of affinity grid assembly. **b**, TEM image at 2500x showing full coverage of GO on grid. Two arrows point to a long wrinkle, which may be due to GO overlapping. Scale bar: 5 µm. **c**, TEM image at 5000x showing GO coating of single layer. Inset indicates selected area electron diffraction (SAED) taken from the center hole marked by dashed square. Scale bar: 1 µm. **d**, Non-cognate tag control grid. A cropped cryo-EM micrograph of human mitochondrial Hsp90 (TRAP1) with SpyTag applied to a SpyTag -PEG (5000Da)-GO grid taken on an Arctica with a K3 camera. Scale bar, 50 nm. **e**, Cropped cryo-EM micrograph of TRAP1-SpyCatcher applied to the same type of Spy-Tag-PEG-GO grid used in d. The image was taken on a Titan Krios with a K2 camera. Scale bar: 50 nm. **f**, Selected 2D averaged classes of nucleotide-bound TRAP1 using affinity grids as described in (e).

To optimize the substrate for coverage and chemical reactivity we synthesized GO using a modified Hummer’s method^15^, rather than using commercially available preparations which have smaller sheets and more variation in oxygen content. Elemental analysis from X-ray photoelectron spectroscopy (XPS) gave a C/O ratio of approximately 2 indicating a relatively high level of oxidation (**Supplementary Fig. 1)**. To reproducibly make EM grids having a uniform GO coverage, we established a simple, robust procedure involving spreading GO films at the air-water interface and lowering them onto submerged grids^16^, derived from a previously described method^17^. We estimated GO coverage at over 90% of grid surface, with ∼40% being monolayer, 40% bilayer and less than 20% having three or more layers. In agreement with our previous experience^16^, we did not notice any negative impact of the GO on contrast in the cryo-EM image even with very small particles. As shown in **Fig. 1b**, a typical TEM image at lower magnification indicates full coverage of GO over the grid square. Monolayer GO (**Fig. 1c**) and double layer GO (**Supplementary Fig. 2**) over carbon holes can be observed by selected area diffraction (insets).

Considering that carboxyl groups mainly decorate on the edges of GO sheets, we decided to use the epoxide groups that cover the bulk of planar area^18^ for functionalization. As illustrated previously^19^, epoxide groups are very efficiently functionalized via nucleophilic reaction with primary amines (**Supplementary Fig. 3**) under non-aqueous conditions. Additionally, under these conditions the much smaller number of carboxyl groups will also react with the amine to form amide bonds. To maximize versatility, we first coupled an amino-PEG-alkyne linker to the grid and then coupled a target-specific reactive azide-PEG spacer in a second step using Cu-free click chemistry^20^. As the GO-PEG-alkyne grids are quite stable, this allows the use of a wide range of coupling chemistries/reactive groups to link the tag catcher to the grid. In the example here, we used an azide-PEG-maleimide to couple to a cysteine on the SpyTag or SpyCatcher to pre-formed PEG-alkyne grids. Alternatively, one could use an azide-PEG-N-hydroxy succinimide to couple to primary amines or directly couple other azide-containing tags. The presence of the PEG segment helps passivate the grid surface, renders it sufficiently hydrophilic to vitrify well, spaces the target away from the surface and also increases the flexibility to minimize potential issues with preferential orientation. We note that proteins did not behave well on functionalized copper grids. We speculate that copper was involved in the Cu-free reaction and released substances which complicated the sample preparation. Holey carbon gold grids were thus used throughout our experiments, although one could just as easily have used gold foil on gold grids.

After coupling the SpyTag or SpyCatcher to the maleimide grids, incubation with a cognate-tagged protein of interest efficiently forms a stable attachment in a matter of minutes. Although we have used this procedure attaching either SpyTag or SpyCatcher to the grid, in the example here we incubated a SpyTag grid with a dilute solution (270nM) of the dimeric mitochondrial Hsp90 molecular chaperone (TRAP1, ∼150KDa) fused to either SpyTag (control) or SpyCatcher (cognate sample, total molecular weight ∼165KDa). The grids were extensively washed to remove unbound or loosely adsorbed proteins. As demonstrated in **Fig 1d**, in the non-cognate control sample, very few TRAP1 molecules are visible. By contrast, applying TRAP1-SpyCatcher to the same SpyTag affinity grids (**Fig. 1e)** results in a very satisfactory particle density and is suitable for high resolution studies. This clearly demonstrates successful and specific affinity capture on the grid with the SpyCatcher/SpyTag system. Obtaining a similar particle density on a regular grid (Quantifoil holey carbon gold grid, 300 mesh) requires a concentration of ∼2 µM, indicating the ability of the affinity grid to specifically concentrate the protein of interest. High-resolution features of TRAP1 are clearly viable in the 2D class averages (**Fig. 1f**) and suggest the data collected are of high quality with minimal impact on contrast from grid modification.

In the current study, we used the human mitochondrial Hsp90 (TRAP1) as a test sample. It is much smaller (∼150 kDa) than samples typically used for testing grid technologies, such as proteasomes and ribosomes, thus making it a stringent test of grid background, contrast, and achievable ice thickness. It also turns out to be particularly sensitive to partial denaturation at the air-water interface (see below). Each TRAP1 protomer within the dimer consists of 3 individual domains: N-terminal domain (NTD), Middle domain (MD), and C-terminal domain (CTD). Together these domains go through a complex ATP binding and hydrolysis cycle^21^ that plays a key role in regulating mitochondrial protein homeostasis and function. Although the crystal structure of TRAP1 from zebrafish had been determined previously^22^, human TRAP1 has proven to be a significant challenge for both X-ray crystallography (producing only poorly diffracting crystals) and cryo-EM. Despite being able to collect a large, high quality dataset, previous attempts of solving the human TRAP1 cryo-EM structure using standard Quantifoil grids produced predominantly a structure where only the TRAP1 NTD and MD from each protomer were resolved (4.1 Å, **Fig. 2a**). Notably, this structure is different from a crystal structure of TRAP1 in which the CTDs were proteolyzed and crystallized post closure^22^. Only a minor population of particles representing full-length TRAP1 could be classified in 3D (class 2 in **Supplementary Fig. 4**), resulting in a medium resolution reconstruction even when starting with a large dataset (4.3 Å, **Fig. 2b**). The dimerization interface between the NTDs of each protomer is preserved in the dominant structure (**Fig. 2a**), which only happens when starting with full length protein. Thus, the disruption of the CTD dimerization interface most likely occurs upon grid preparation, presumably due to the interaction with the air-water interface, resulting in a flexible or denatured CTD invisible in our reconstructions.

**Figure 2:**
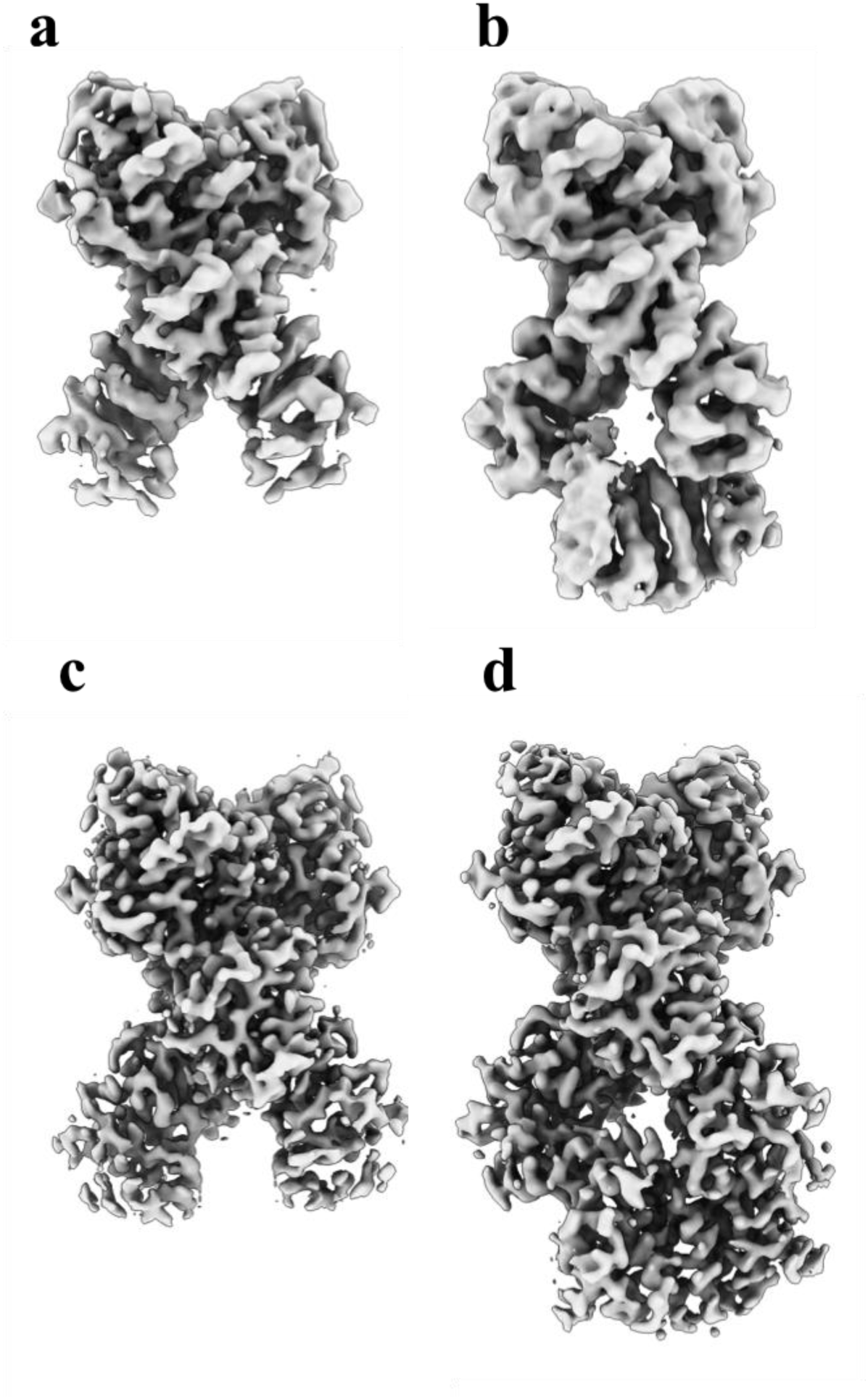
3D density map of human TRAP1 structures determined using both regular Quantifoil grids and GO based affinity grids. a, A TRAP1 structure (4.1 Å) with flexible/disrupted CTD obtained from the dominant class using regular Quantifoil grid. b, A full-length TRAP1 structure (4.3 Å) obtained from a minor class of particles using regular Quantifoil grids. c, Structure of TRAP1 with SpyCatcher with flexible CTD at 3.1 Å resolution from a minor class using GO based affinity grid. d, Full length structure of TRAP1 with SpyCatcher obtained from am more dominant class using GO based affinity grid at 3.3 Å resolution.

Fortunately, our GO based affinity grids solve this problem, and full-length particles are easily visible in the raw micrographs (**Fig. 1e**) as well as in the 2D class averages (**Fig. 1f)**. We hypothesize that the preservation of the TRAP1 CTD dimerization interface is due to the affinity tags keeping the protein away from the air-water interface. As demonstrated by tomography, the PEG spacer keeps the protein away from both the air-water interface and the GO surface, thereby avoiding potentially unfavorable interactions with regions of unmodified GO (**Supplementary Fig. 5**). Although the TRAP1 dimer state missing the CTD density is still present in our affinity grid dataset (reconstructed to 3.1 Å resolution, Fig 2c), the ratio between full-length TRAP1 population and CTD missing population was improved from 1:6.5 to 1:2.3 after discarding low quality particles through 3D classification (**Supplementary Fig. 4 and 6**). Using the affinity grids, we were able to solve the full-length TRAP1 structure at 3.3 Å resolution (**Fig. 2c**). This structure closely resembles the crystal structure of zebrafish TRAP1 and exhibits the same pronounced asymmetry^22^, proving that this distinctive conformation is both conserved across species and that it is not a consequence of crystallization. Having datasets for the same sample using both conventional grids and the affinity grids reveals clear differences in orientation bias. From the 2D averages, side views are quite rare with conventional grids, but prevalent with the affinity grids. The angular distributions for each final reconstruction are shown in **Supplementary Fig. 7**.

To further evaluate the SpyCatcher/SpyTag affinity grid, we also performed testing in the reverse manner, immobilizing SpyCatcher on the grid, and then applying TRAP1-SpyTag. We also explored two different PEG chain lengths (600Da vs 5000 Da M.W.) in the azide-PEG-maleimide spacer. While we did not pursue full reconstructions in these cases, there were no apparent differences in either particle density or 2D class average quality (**Supplementary Fig. 8-16**).

In summary, our GO based affinity grids provide selective enrichment of a tagged sample on the grid without negatively impacting image contrast and particle orientation. This allows even small particles to be readily reconstructed at high-resolution. Perhaps even more importantly, the ability of our affinity grids to protect delicate samples from partial denaturation/aggregation at the air-water interface is a critical and enabling technology. Here this allowed us to determine the full-length structure of human TRAP1 at atomic resolution for the first time, paving the way for future structure-based drug discovery experiments.

It is noteworthy that our choice of grid modification strategies readily allows the use of other affinity tagging strategies beyond the SpyCatcher/SpyTag system employed here. For example, HaloTag ligand can be coupled to the grids allowing the purification of HaloTagged proteins, often favored by cell biologists for the ability to incorporate small molecule fluorophores for in vivo light microscopy^23^. DNA, nanobodies, etc. can also be readily incorporated, extending the practical applications to a much broader range of biological problems. Alternatively, the grid surface can be modified to display primary or secondary amines by functionalization with diamine species, or other chemical chemotypes. For example, we have found that amino-GO grids wet better, promote protein adsorption and improve orientation distribution compared with bare GO grids. Exploring these variants is part of our on-going research and will be discussed in a separate paper.

## ACKNOWLEDGEMENTS

We wish to thank Eugene Palovcak for helpful discussions, Michael Braunfeld, Alex Myasnikov, David Bulkley for help and running the UCSF Advanced Cryo-Electron Microscopy Facility, and Matt Harrington for HPC support. Funding to DAA comes from a UCSF PBBR technology development grant, and NIH grants R35GM118099, U54 CA209891, U01MH115747, U19AI135990. YC is supported by NIH grants R01GM098672, P50GM082250, P01GM11126. DAA and YC are supported by the Howard Hughes Medical Institute, and the facility has received NIH instrumentation grants S10OD020054 and S10OD021741. YL is supported by the HHMI-Helen Hay Whitney Foundation Postdoctoral Fellowship and the American Heart Association Postdoctoral Fellowship.

## CONTRIBUTIONS

F.W. and D.A.A. designed the experiments. F.W. developed affinity grids and implemented protein vitrification. Y.L. collected and analyzed the single particle cryoEM data. Z.Y. purified protein and contributed to data collection. S.L. and Z.Y. performed tomography analysis. F.W., Y.L. and Z.Y. wrote the manuscript with input from all authors. Y.C. and D.A.A. supervised the project.

## CORRESPONDING AUTHOR

Correspondence to David A. Agard

## ONLINE METHODS

### GO synthesis

GO was synthesized via the modified route developed by Marcano et al.^15^. Into the mixture of concentrated sulfuric acid (120 ml, Ward’s Science 470302-872) and phosphoric acid (13 ml, Sigma-Aldrich 345245), graphite flakes (1 g, Sigma-Aldrich 332461) were added. Potassium permanganate (6 g, Sigma-Aldrich Sigma-Aldrich 223468) was then added very slowly into the system before it was moved into a water bath of 45 °C and stirred overnight. After the reaction was moved into an ice bath, DI water (100 ml) was poured in followed by addition of 30% H2O2 (1.5 ml, Sigma-Aldrich 216763). The mixture was allowed to sit for 2 hours and then centrifuged (5000 rpm, 20 min). The solid material at the bottom was retrieved and washed extensively with DI water by centrifugation until the pH reached 5. Finally, the remaining viscous material was collected and stirred overnight to make a GO stock solution in water. After drying at 80 °C, XPS measurement (PHI 5600) was performed on GO flakes.

### GO deposition onto EM grids

To coat GO sheets onto EM grids, we revised the method of Langmuir-Blodgett assembly which was described by Cote et al.^17^, and also reported in our previous work^16^. The GO water stock solution was diluted with methanol/water (5:1, v:v) to a concentration of 0.1 mg/ml. Mild stirring for 30 mins rather than sonication was used to avoid destruction of GO sheets, producing a GO working solution. An epoxy coated stainless steel mesh (Mcmaster-Carr) stand was placed at the bottom of a glass petri dish (60 mm in diameter, 15 mm in height) and DI water was filled to the top. EM grids (Au Quantifoil, 300 mesh) were used as received and placed on the mesh with carbon side facing upward. Then a total volume of 230 µl of GO working solution was spread dropwise onto the water surface at different spots at a speed of 50 µl/min using a syringe. After the water was drained, the GO coated grids were dried at room temperature overnight for use. Coverage of GO was examined on a Tecnai 20 TEM (Thermal Fisher Scientific) with an acceleration voltage of 200 kV.

### Surface modification of GO coated EM grids

In a 1.5 ml centrifuge microtube, one GO coated EM grid (GO grids) was submerged in 20 µl of dibenzocyclooctyne-polyethylene glycol-amine (DBCO-PEG4-Amine, Click Chemistry Tools A103P) solution in dimethyl sulfoxide (DMSO) at a concentration of 10 mM and shaken at room temperature overnight. Following that, the DBCO functionalized grid was washed by DMSO and DI water for three times, respectively, and submerged in 20 ul of azide-polyethylene glycol-maleimide (M.W. 600 Da, Nanocs PG2-AZML-600; or M.W. 5000 Da, Nanocs PG2-AZML-5k) solution in DI water. The reaction was shaken at room for 6 hours and the grid was washed by DI water and ethanol for three times, respectively. The as-made maleimide grid was then dried in ambient for half an hour and stored at −20°C for future use.

### Protein purification

The constructs for human TRAP1 (hsTRAP1) fused either SpyTag or Spy Catchter were the same as previously used^24^. Proteins were expressed in *E. coli* BL21-star (DE3) cells. Cells were grown overnight at 18 °C after induction with 0.5 mM isopropyl-β-D-thiogalactopyranoside (IPTG) at OD600 ∼ 0.7. Cells were harvested by centrifugation (3,500 g for 20 min), resuspended in buffer A (25 mM HEPES pH 7.5, 150 mM NaCl, 10mM Imidazole and 5% glycerol), then lysed by Emulsiflex. The lysate was clarified by centrifugation (27,000 g for 30 min) then subjected to a prepacked 5-ml His-trap column (GE Healthcare) for binding. After 50ml buffer A wash, Buffer B (25 mM HEPES pH 7.5, 150 mM NaCl, 280 mM Imidazole and 5% glycerol) was used to elute off the protein from the His-trap column. The protein was concentrated in a 30,000 MWCO concentrator (Amicon) for further purification on a Superose 6 size-exclusion column (GE Healthcare) equilibrated in buffer C (25 mM HEPES pH 7.5, 150 mM NaCl and 5% glycerol).

### Assembly and vitrification using affinity grids

A buffer with a pH of 7.5 containing 20 mM HEPES, 60 mM KCl was used throughout the process. SpyCatcher was diluted with buffer to a final concentration of 1.4 µM. SpyTag with a sequence of VPTIVMVDAYKRYKC was purchased from Biomatik. SpyTag was first dissolved in DMSO to a concentration of 560 µM and then diluted with buffer to a final concentration of 2.8 µM. TRAP1 with SpyCatcher or SpyTag was diluted in buffer with 1 mM ADP-BeF_x_/MgCl_2_ and was incubated at 37 °C for 30 min to achieve a homogenous closed state of TRAP1^24^. The concentration of TRAP1 was 270 nM in all cases.

For affinity testing with SpyTag on grid, one maleimide grid was first incubated in 200 ul of SpyTag solution at room temperature for 2 hours. The grid was washed with DI water and ethanol 3 times, and dried in ambient for 1 hour. Then the grid was picked up by tweezers from Vitrobot IV and placed horizontally. In ambient, 3 ul of solution of TRAP1 with SpyCatcher was applied to carbon side of the grid and incubated for 4 min. The grid was washed by flipping the grid downward and touching 30 ul of buffer droplet with carbon side on parafilm for three times. After each wash, liquid was drained by filter paper. Then 3 ul of solution of TRAP1 with SpyCatcher was applied to carbon side of the grid and incubated for another 4 min. The grid was washed with buffer using the same way mentioned above. Very quickly 3 ul of buffer was applied to the grid before it was completely dried. Tweezers with the grid was retracted to Vitrobot chamber and blotted immediately using the following condition: 22 °C, 100% humidity, blot force 1, blot time 8 s. Grid was plunged into liquid ethane and stored in liquid nitrogen. For the negative control, TRAP1 with SpyTag was applied instead to the grids.

### Cryo-EM data acquisition

A total of 5 cryo-EM datasets ware collected using either Titan Krios or Talos Arctica (Thermo Fisher Scientific). For the first dataset, TRAP1 fused with SpyCatcher are applied to GO grids functionalized with SpyTag and PEG spacer M.W. 5000 Da. A total of 2670 micrographs were collected using beam-image shift^25^ on a Titan Krios microscope (Thermo Fisher Scientific) operated at 300 kV with a K2 Summit direct electron detector (Gatan, Inc.) and a slit width of 20 eV on a GIF-Quantum energy filter. Images were recorded with SerialEM in super-resolution mode with a pixel size of 0.407 Å. Defocus varied from 0.8 um to 2.5 um. Each image was fractionated to 80 frames (0.1 sec each, total exposure of 8 seconds) with dose rate of 8.6 e/Å^2^/sec for a total dose of 68.8 e/Å^2^.

The other 4 datasets were collected on a Talos Arctica (Thermo Fisher Scientific) equipped with a K3 detector (Gatan, Inc.). The microscope was operated at 200 kV at a nominal magnification of either 36,000x or 28000x, corresponding to super-resolution pixel sizes of 0.57 Å/pix or 0.72 Å/pix. Defocus varied from 0.8 um to 2.5 um. Each image was fractionated to 100 frames (0.03 sec each, total exposure of 3 seconds) with dose rate of 24.7 e/Å^2^/sec for a total dose of 74.1 e/Å^2^. For more details see **Table S1**.

### Image processing

Image stacks were motion corrected and summed using MotionCor2^26^, resulting in Fourier-cropped summed images (0.814 Å/pix for Krios1 data, 1.14 Å/pix for Arctica data at 36000x, and 1.08 Å/pix for Arctica data at 28000x). CTFFIND4 was used to estimate defocus parameters for all the images^27^. Initial particle picking was carried out using Gautomatch without a template to generate the 2D class averages, which were then used as templates for a second-round particle picking on micrographs with 25 Å low-pass filtering. Relion 3.0 beta was used for all the following steps. Reference free 2D classification were performed for 25 iterations using images binned by 4. Bad particles were excluded based on 2D average results. The remaining particles were 3D classified into 10 classes using images binned by 2 and an initial model generated from the crystal structure of zebrafish TRAP1 and lowpass filtered to 50 Å. The particles from the 3D classes with high-resolution features were reextracted without binning and subjected to 3D auto-Refinement in Relion. The resulting maps from refinement were post-processed and sharpened by an automatically estimated B-factor in RELION. All resolutions were estimated by applying a soft mask around the protein density and the gold-standard Fourier shell correlation (FSC) = 0.143 criterion.

### Tomogrphy analysis

The electron cryo-tomography tilt series of TRAP were collected on a Talos Arctica microscope running at 200 KV by SerialEM^28^. A K3 camera was used for recording movies in the counting mode. The pixel size on micrograph is 2.60 Å. Data were acquired in the tilt range ±60° at 3° interval, in two branches starting at 0°, at 2-4 µm underfocus with a cumulative dose of 60 *e*^-^/Å^2^. The tilt series’ were aligned by IMOD^29^ based on the gold fiducials embedded with the protein sample. Tomograms were reconstruted by TOMO3D^30^.

