## Supplementary Information for "General and robust covalently linked graphene oxide affinity grids for high-resolution cryo-EM"

Department of Biochemistry & Biophysics and the Howard Hughes Medical Institute. University of California, San Francisco. San Francisco, CA 94143

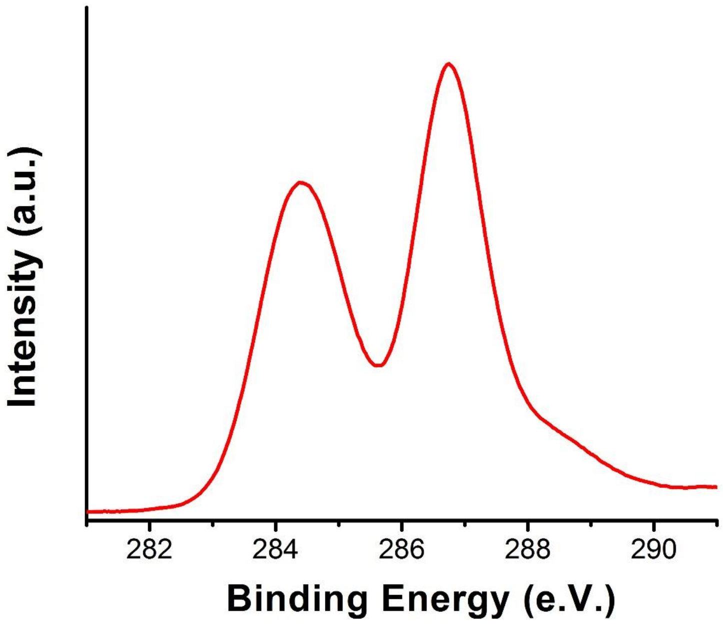


**Supplementary Figure 1**: C1s XPS spectrum of GO synthesized.

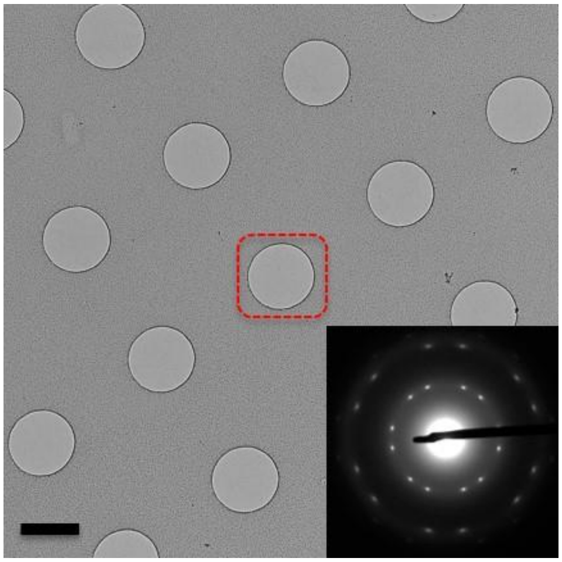

**Supplementary Figure 2**: TEM image at 5000x showing GO coating of double layers on EM grids. Inset indicates selected area electron diffraction (SAED) pattern taken from the center hole marked by dash square. Scale bar: 1 μm.

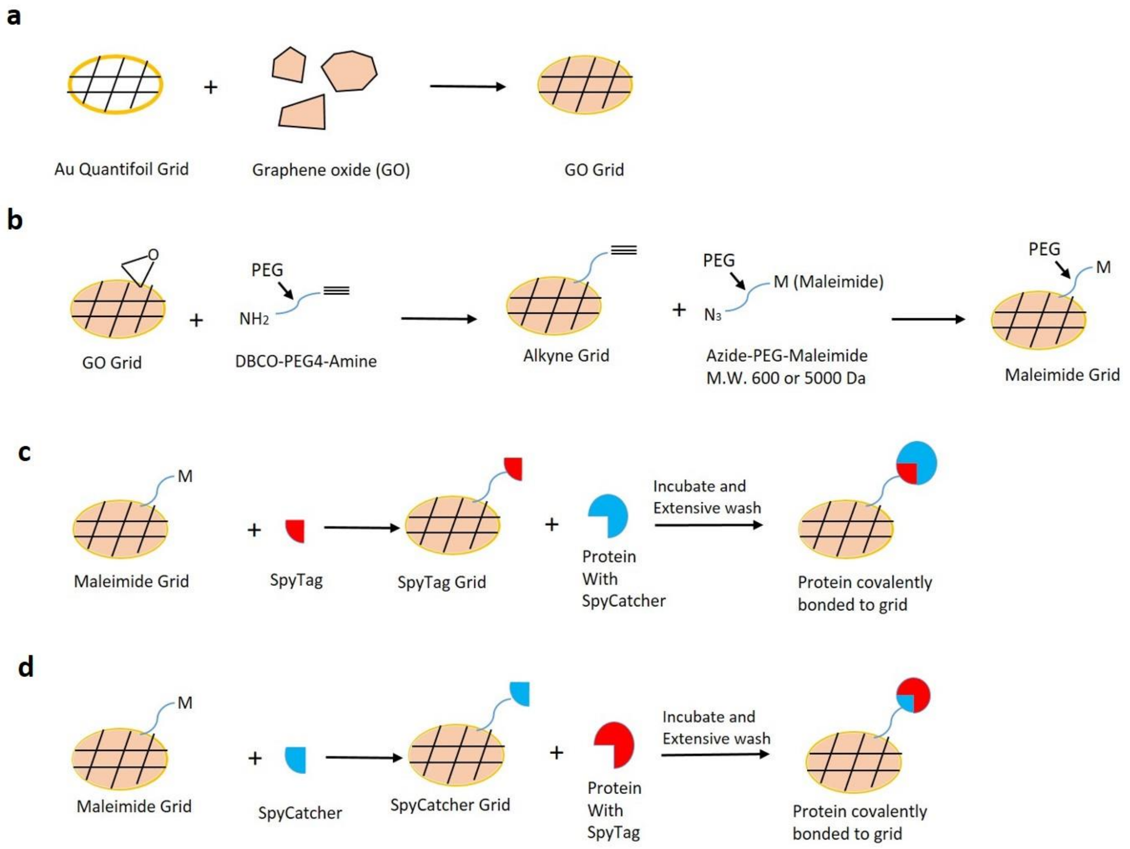


**Supplementary Figure 3**: Schematic illustration of affinity grid assembly and testing routes. **a**, GO coating onto EM grids. **b**, Surface functionalization of GO coated grids. **c**,**d**, Assembly and testing of affinity grids. In **c**, SpyTag is deposited on the grid and SpyCatcher is fused onto protein. In **d**, SpyCatcher is deposited on the grid and SpyTag is fused onto protein.

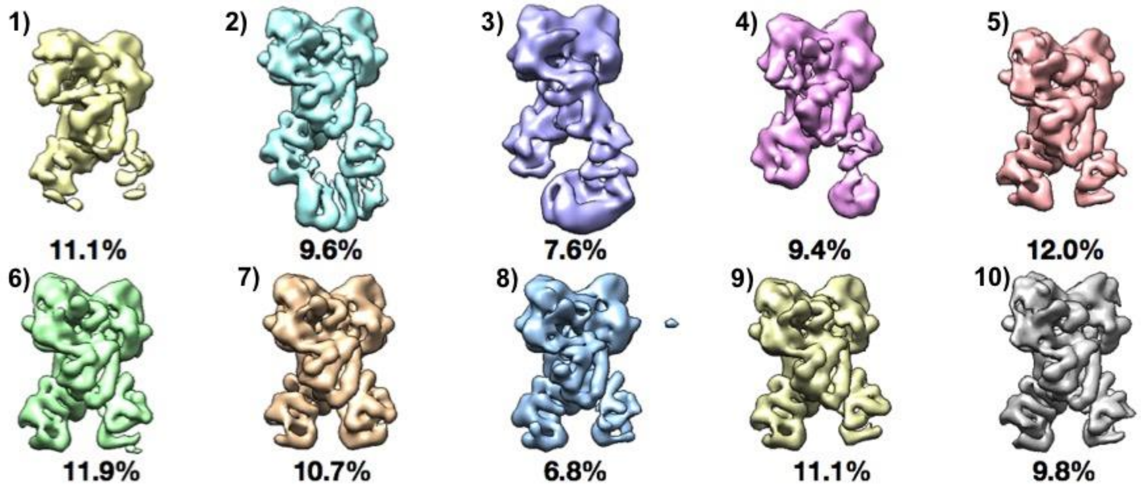


**Supplementary Figure 4**: The final round of 3D classification of TRAP1 with SpyCatcher on regular Quantifoil grids. Classes 5-10 represented the TRAP1 dimer with missing CTD density. Class 10 was further refined to 4.1 Å as shown in Figure 2a. Class 2 represented the full-length TRAP1 dimer and was further refined to 4.3 Å as shown in Figure 2b. Classes 1, 3, 4 were discarded as low-quality particles.

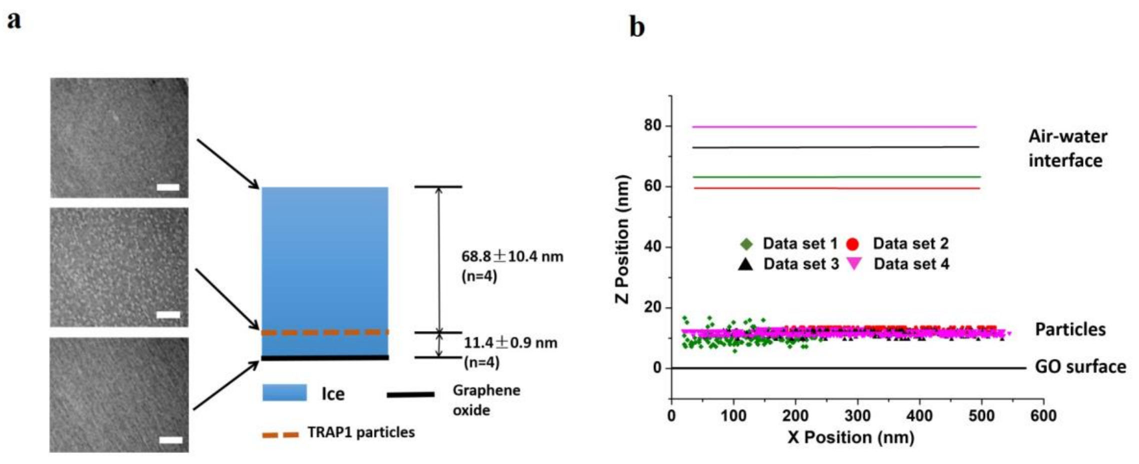


**Supplementary Figure 5**: Determination of particle position from within the ice layer by tomographic analysis. SpyTag was anchored to GO grid via a PEG spacer (M.W. 5000 Da) and SpyCatcher was fused to TRAP1 protein. **a**, Images correspond to three representative sections along Z-axis from the tomography reconstruction. In the example on the left, TRAP1 particles are approximately 69 nm away from the air-water interface and 11.4 nm from the GO surface. Scale bar: 50 nm. **b**, Localization of TRAP1 particles in 4 data sets (represented by markers in different colors) in (a). GO surface is set to zero for all data sets. Air-water interfaces are illustrated with lines of color corresponding to their own data set.

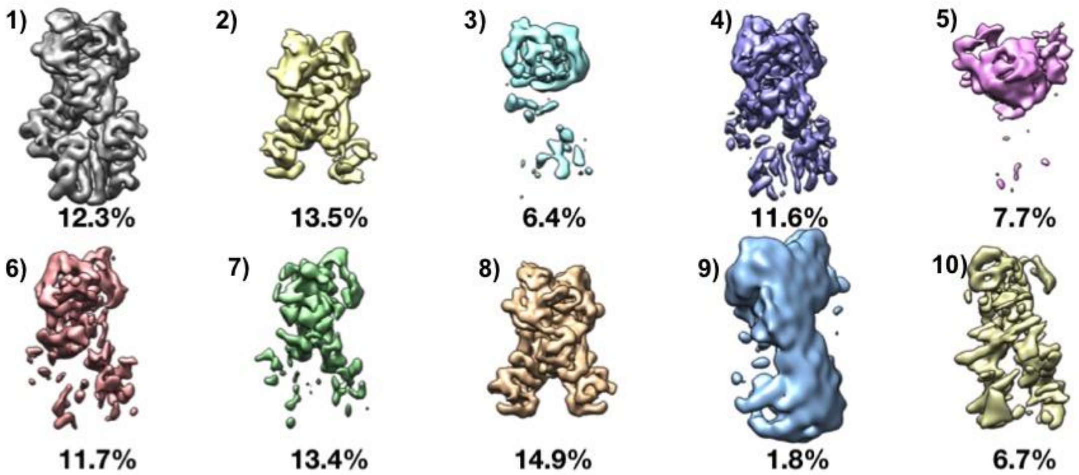


**Supplementary Figure 6**: 3D classification of TRAP1 fused with SpyCatcher on GO based affinity grid. Class 1 represented the full-length TRAP1 dimer and were further refined to a resolution of 3.3 Å as shown in Figure 2d. Classes 2 and 8 represented TRAP1 dimer with missing CTD density. Class 8 was further refined to a resolution 3.1 Å as shown in Figure 2c. The other 7 classes were discarded as low-quality particles.

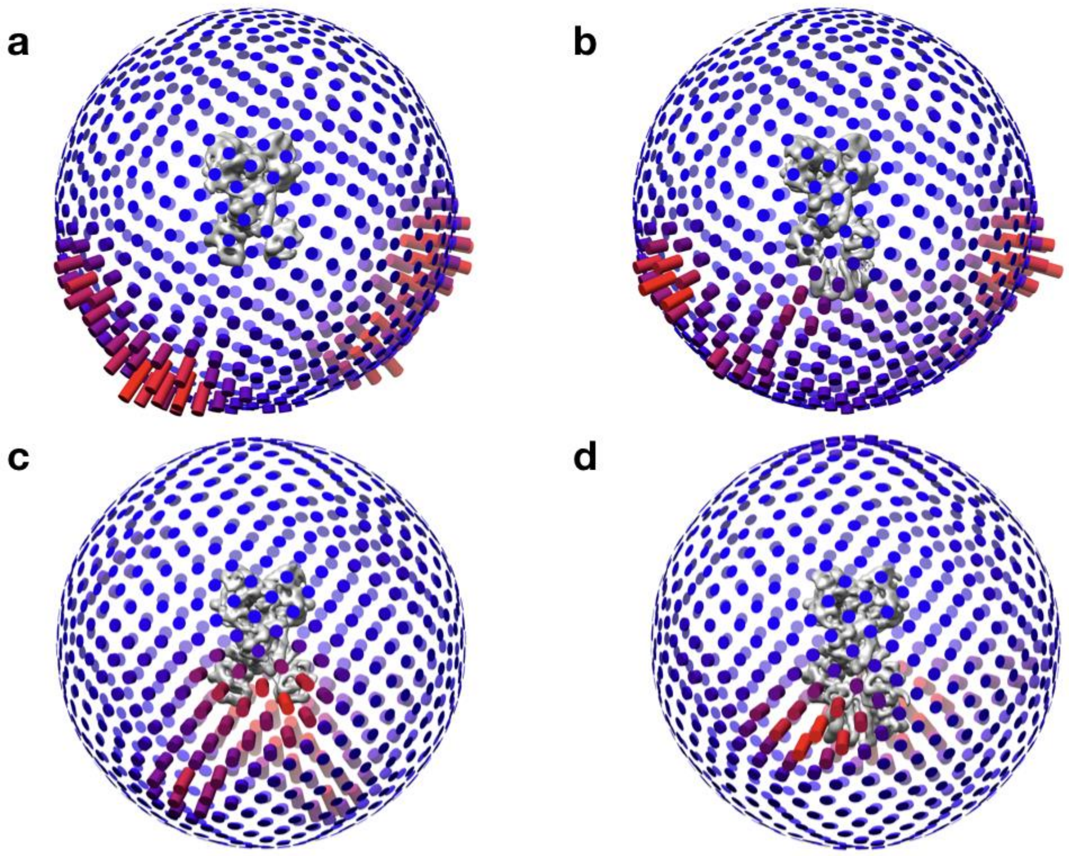


**Supplementary Figure 7**: Orientation distribution of TRAP1 particles using Quantifoil grid and GO based affinity grid for structures shown in Fig. 2. The orders in the two figures are the same.

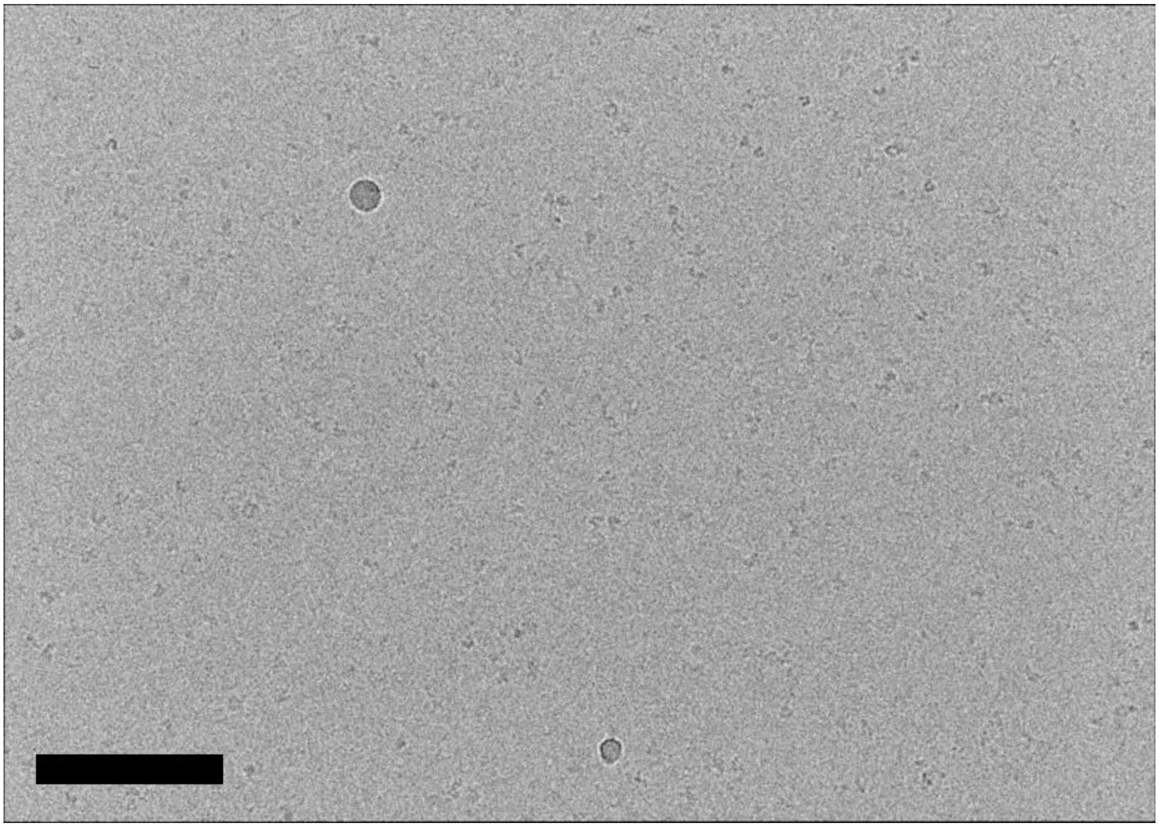

**Supplementary Figure 8**: Cryo-EM micrograph of TRAP1 with SpyTag in negative control group. SpyTag was attached to grid via PEG spacer with M.W. of 600 Da. Scale bar: 100 nm.

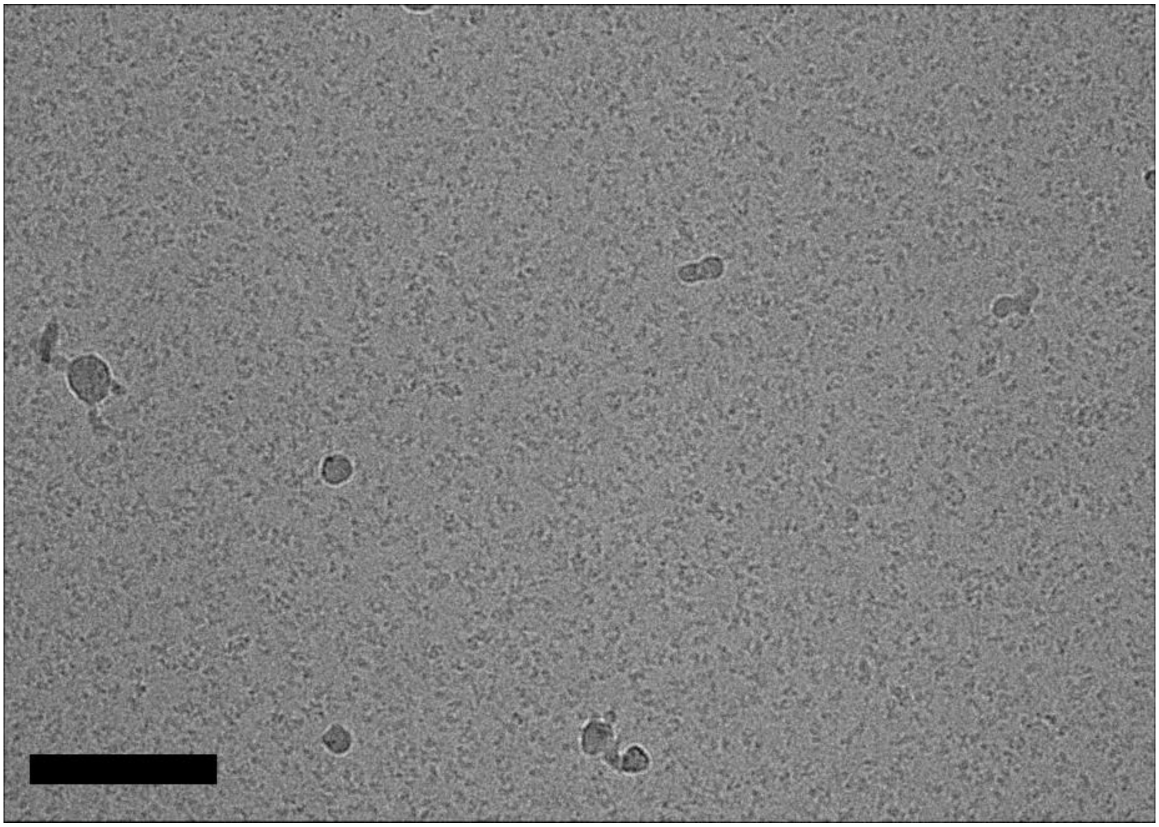

**Supplementary Figure 9**: Cryo-EM micrograph of TRAP1-SpyCatcher bound to a SpyTag-PEG spacer (600 Da) affinity grid. Scale bar: 100 nm.

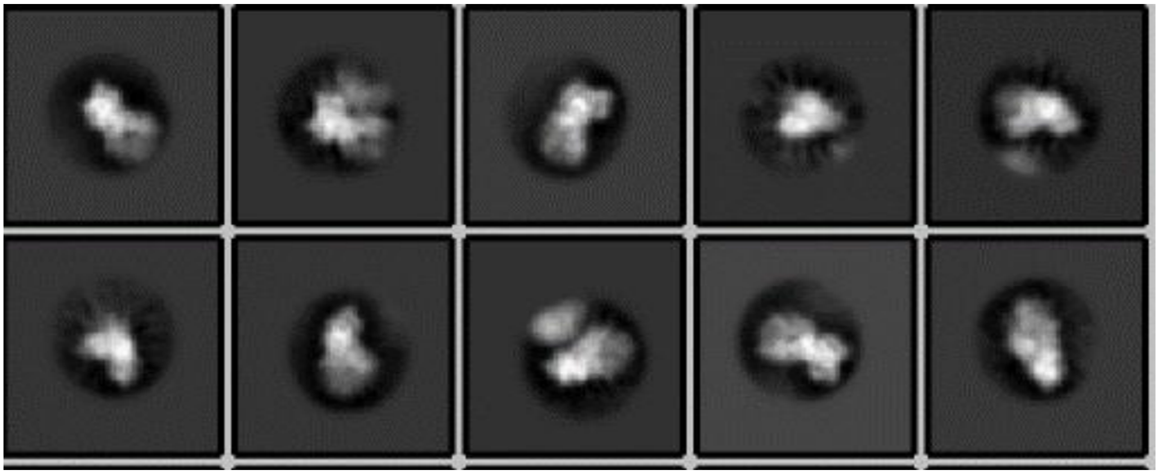

**Supplementary Figure 10**: 2D class averages from images of TRAP1:SpyCatcher on affinity grid as in Supplementary figure 9

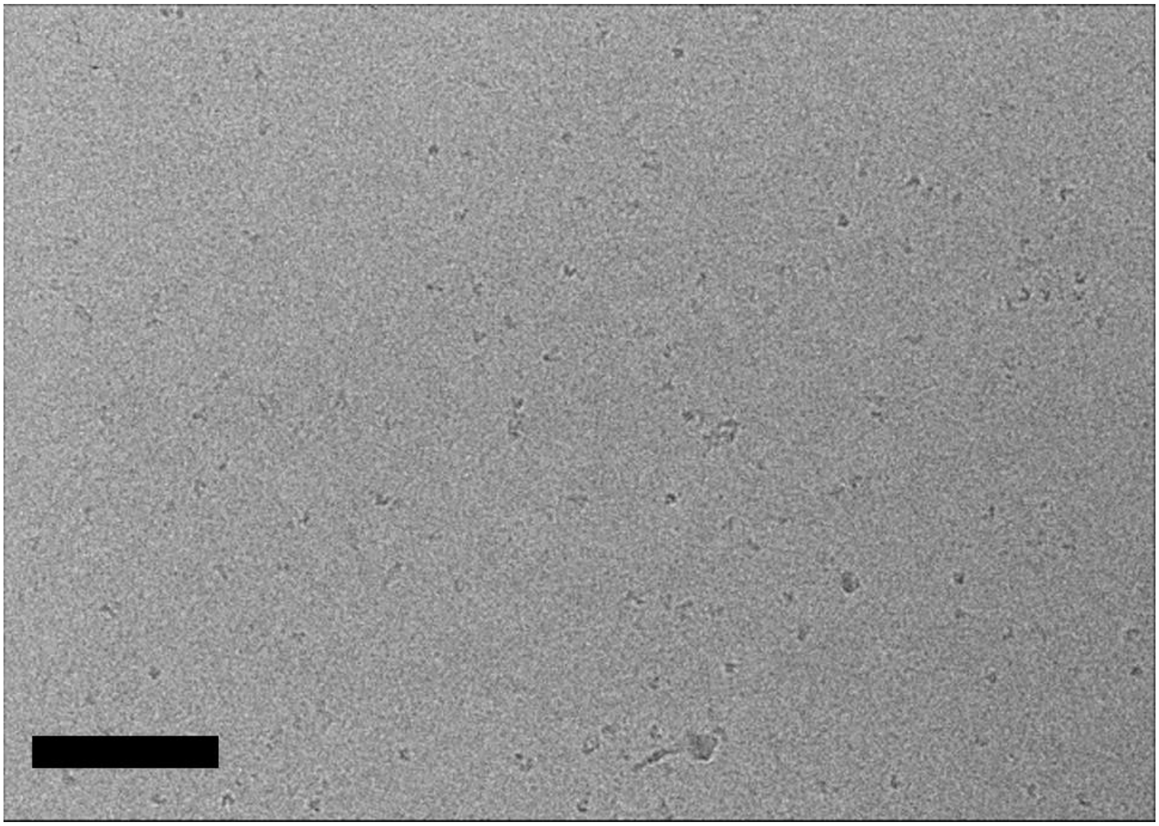

**Supplementary Figure 11**: Cryo-EM micrograph of TRAP1:SpyCatcher incubated with SpyCatcher affinity grid as a negative control. PEG spacer had M.W. of 5000 Da. Scale bar: 100 nm.

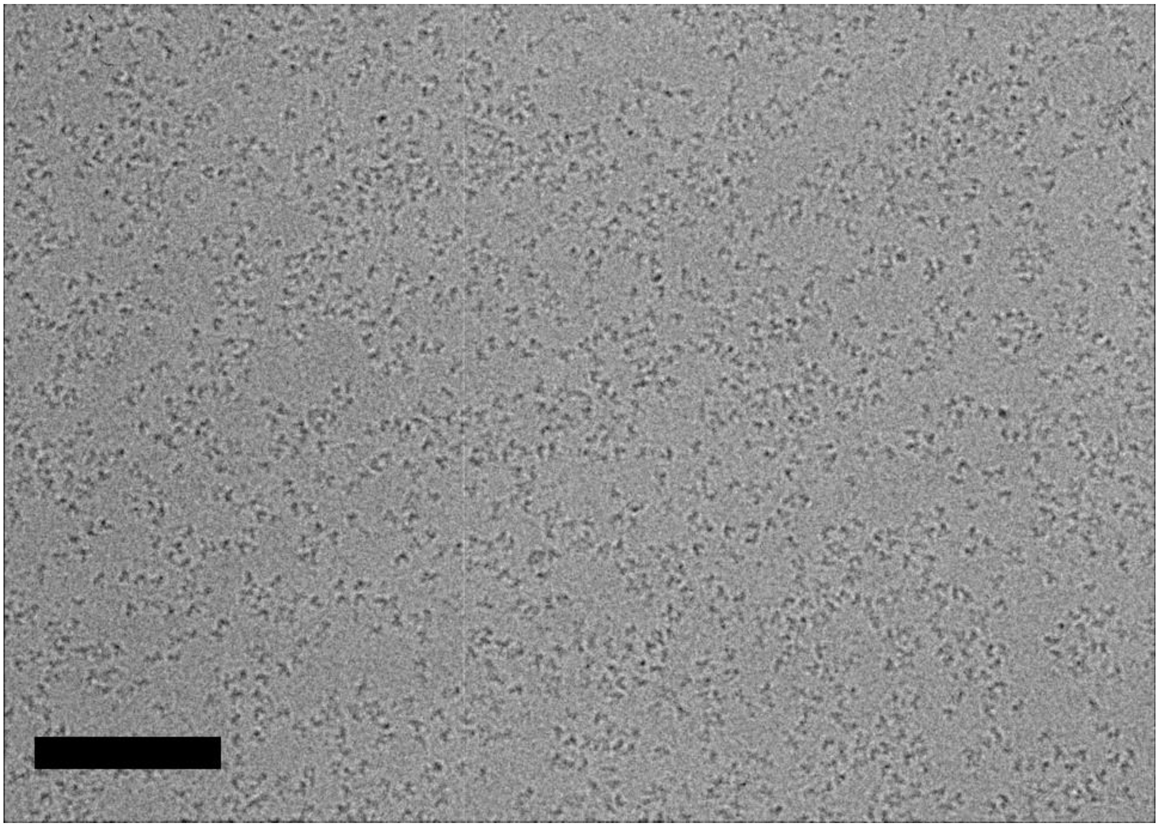

**Supplementary Figure 12**: Cryo-EM micrograph of TRAP1:SpyTag with SpyCatcher affinity grid PEG spacer had M.W. of 5000 Da. Scale bar: 100 nm.

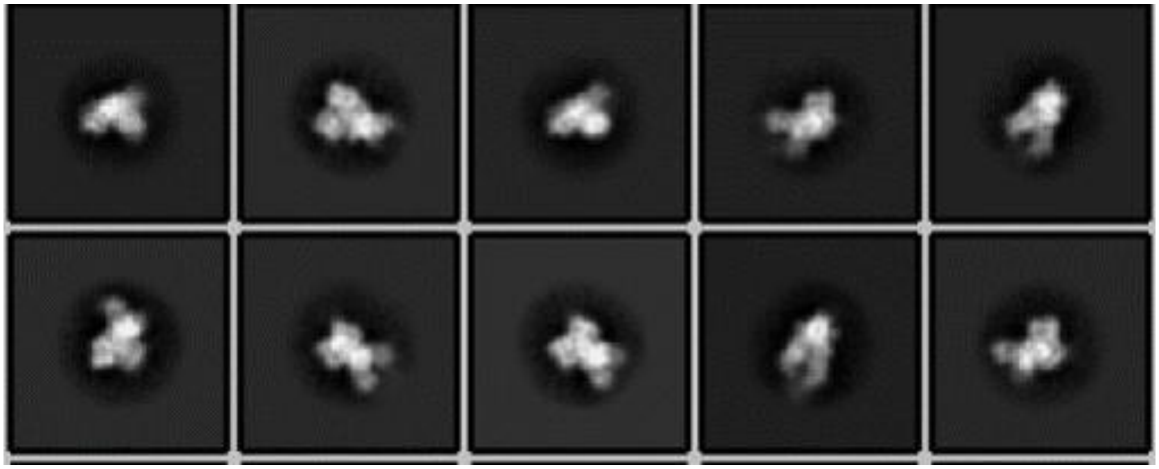

**Supplementary Figure 13**: 2D class averages of TRAP1:SpyTag with SpyCatcher grid. PEG spacer had M.W. of 5000 Da.

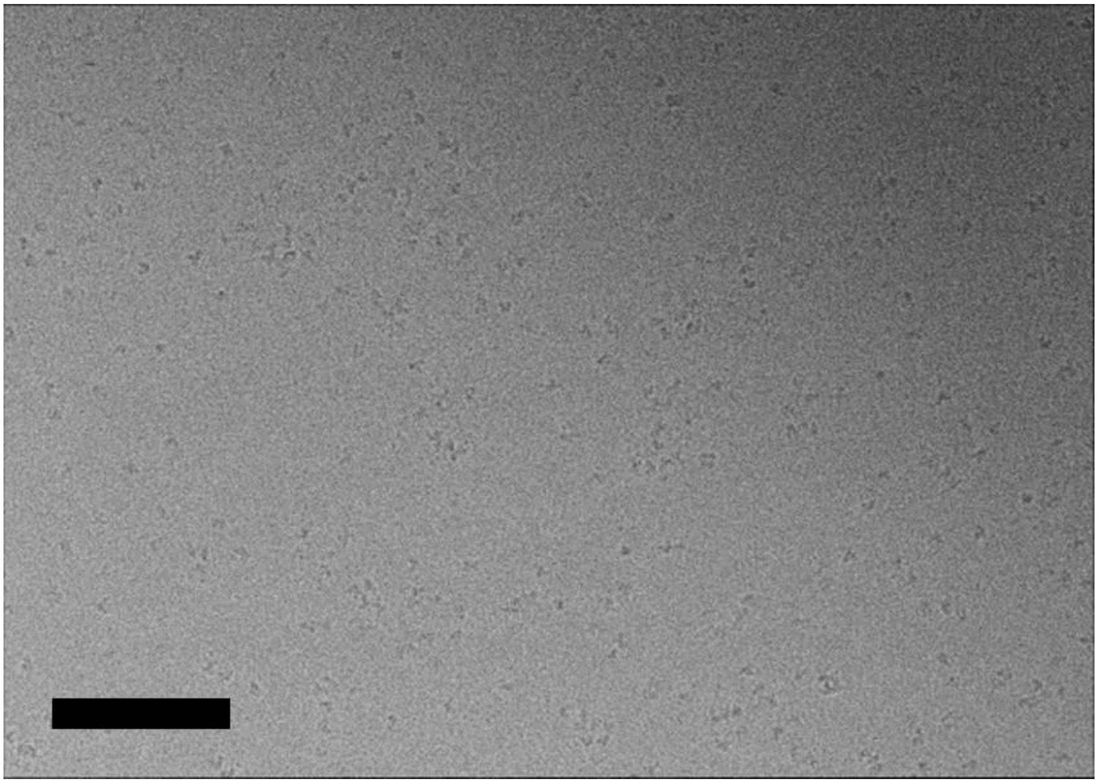

**Supplementary Figure 14**: Cryo-EM micrograph of TRAP1:SpyCatcher interacting with SpyCatcher affinity grid as a negative control. PEG spacer had M.W. of 600 Da. Scale bar: 100 nm.

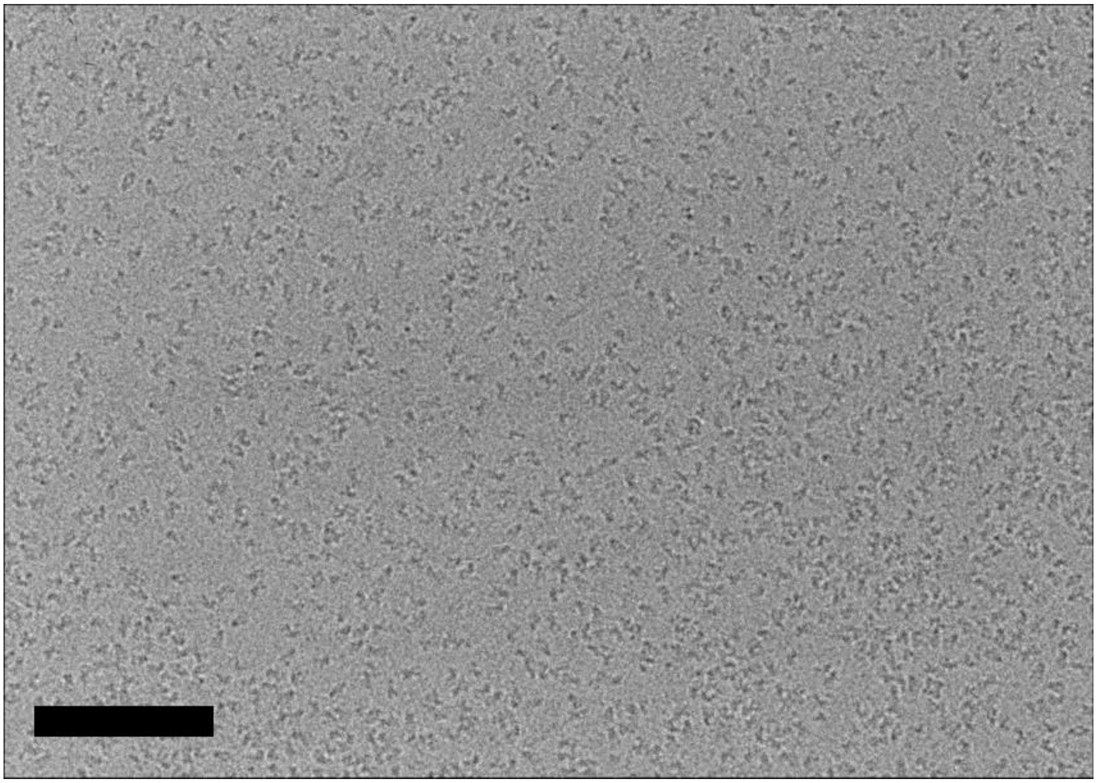

**Supplementary Figure 15**: Cryo-EM micrograph of TRAP1:SpyTag interacting with SpyCatcher affinity grid. PEG spacer had M.W. of 600 Da. Scale bar: 100 nm.

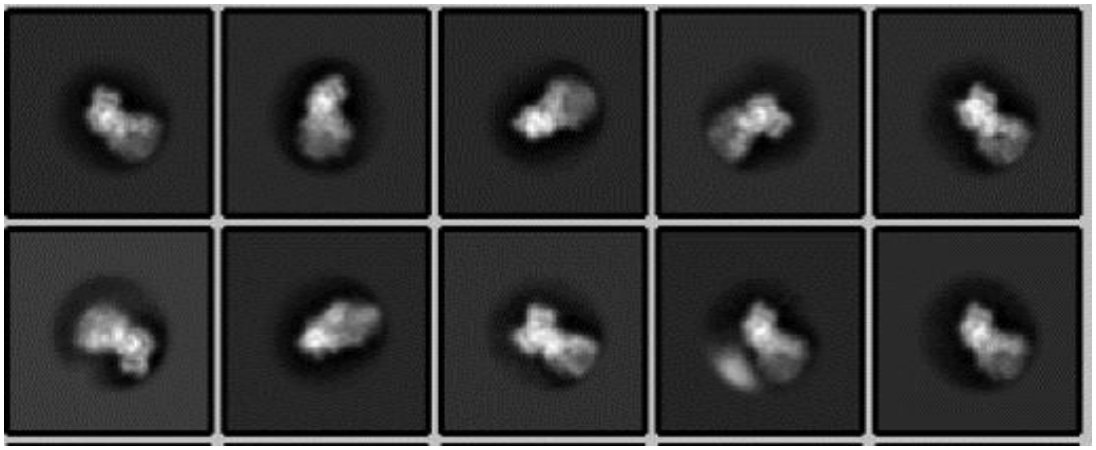

**Supplementary Figure 16**: 2D class averages of TRAP1:SpyTag with SpyCatcher affinity grid, 600 Da PEG spacer.

**Table S1. Summary of collected data**

| Sample | TRAP1 SpyCacther | TRAP1 SpyCacther | TRAP1 SpyCacther | TRAP1 SpyTag | TRAP1 Spytag |
|---|---|---|---|---|---|
| Grid | GO affinity With SpyTag | Regular Quantifoil | GO affinity With SpyTag | GO affinity With SpyCatcher | GO affinity With SpyCatcher |
| PEG spacer M.W. | 5000 | n/a | 600 | 5000 | 600 |
| Microscope | Titan Krios | Talos Arctica | Talos Arctica | Talos Arctica | Talos Arctica |
| Voltage | 300 kV | 200 kV | 200 kV | 200 kV | 200 kV |
| Energy filter | Yes | No | No | No | No |
| Camera | K2 | K3 | K3 | K3 | K3 |
| Pixel size (A/pix) | 0.814 | 1.14 | 1.14 | 1.08 | 1.14 |
| micrographs | 2667 | 1984 | 141 | 234 | 209 |
